# LotOfCells: data visualization and statistics of single cell metadata

**DOI:** 10.1101/2024.05.23.595582

**Authors:** Óscar González-Velasco

## Abstract

Single-cell sequencing unveils a treasure trove into the biological and molecular characteristics of samples. Yet, within this flood of data, the challenge to draw meaningful conclusions sometimes can be time consuming and a tortuous process.

Here we introduce LotOfCells: a simple R package designed to explore the intricate landscape of phenotypic data within single-cell studies. Normally, we are interested in visualizing and measuring if the differences in the proportion of number of cells across various covariates is significant or biologically relevant. As an example, one of the most common questions is the proportion of different cell types across conditions in our experiment, or the cluster composition before and after treatment (e.g.: difference in cell type proportions between wild type and mutant). LotOfCells helps with the interpretation and visualization of meta-data of these recurrent scenarios, including the test of proportion changes across multiple ordered stages. Additionally, it computes a symmetric divergence score to measure global deregulation of cell proportions due to a condition.

**Code repository:** R package, manual and relevant examples can be accessed on the GitHub repository: *https://github.com/OscarGVelasco/LotOfCells*

## 1 Introduction

Single-cell sequencing has revolutionized our ability to study cellular heterogeneity, offering insights into diverse biological processes. Analyzing single cell data requires sophisticated computational methods to decipher complex patterns and identify meaningful differences between cell populations [1].

Once the sequencing data has been pre-processed and aligned, an archetypal single cell data analysis pipeline usually starts by visualising and exercising an exploratory analysis [2–4]. Single cell data possess numerous challenges due to the high-dimensional nature of the data, sparsity, and technical noise [5]. At this initial state of the analysis, visualisation methods are critical to assess the quality, content, and heterogeneity of the dataset [6–8].

There are numerous software tools and packages for analysing and visualising single cell data [6, 8–19], therefore, here is another one: variety is the spice of life.

A critical aspect of single cell analysis is comparing cell type proportions or other covariates (e.g.: cluster or tissue composition) between different experimental conditions, disease states, or biological samples. Conventional statistical tests may not effectively capture subtle differences in covariate distributions, and assessing how extreme and significance our observations are can be challenging due to the limited power of classical statistical approaches, necessitating the development of more sensitive and robust methods.

Here we present LotOfCells, an R package to easily visualize and analyze the phenotype data (metadata) from single cell studies. It allows to test whether the proportion of the number of cells from a specific population is significantly different due to a condition or covariate (e.g.: number of cells per cell type changes between tumor and control). The numerous options for visualizing the data comes with ready-to-publish figures. The package is compatible with Seurat and SingleCellExperiment objects.

## 2 Results

We used LotOfCells to test and visualise a public scRNA-seq dataset from metastatic lung adenocarcinoma from Kim et al [20]. Composed by 208,506 cells, and including normal adjacent tissues, and early to metastatic stage cancer across five different stages, in 44 patients.

**Fig. 1:**
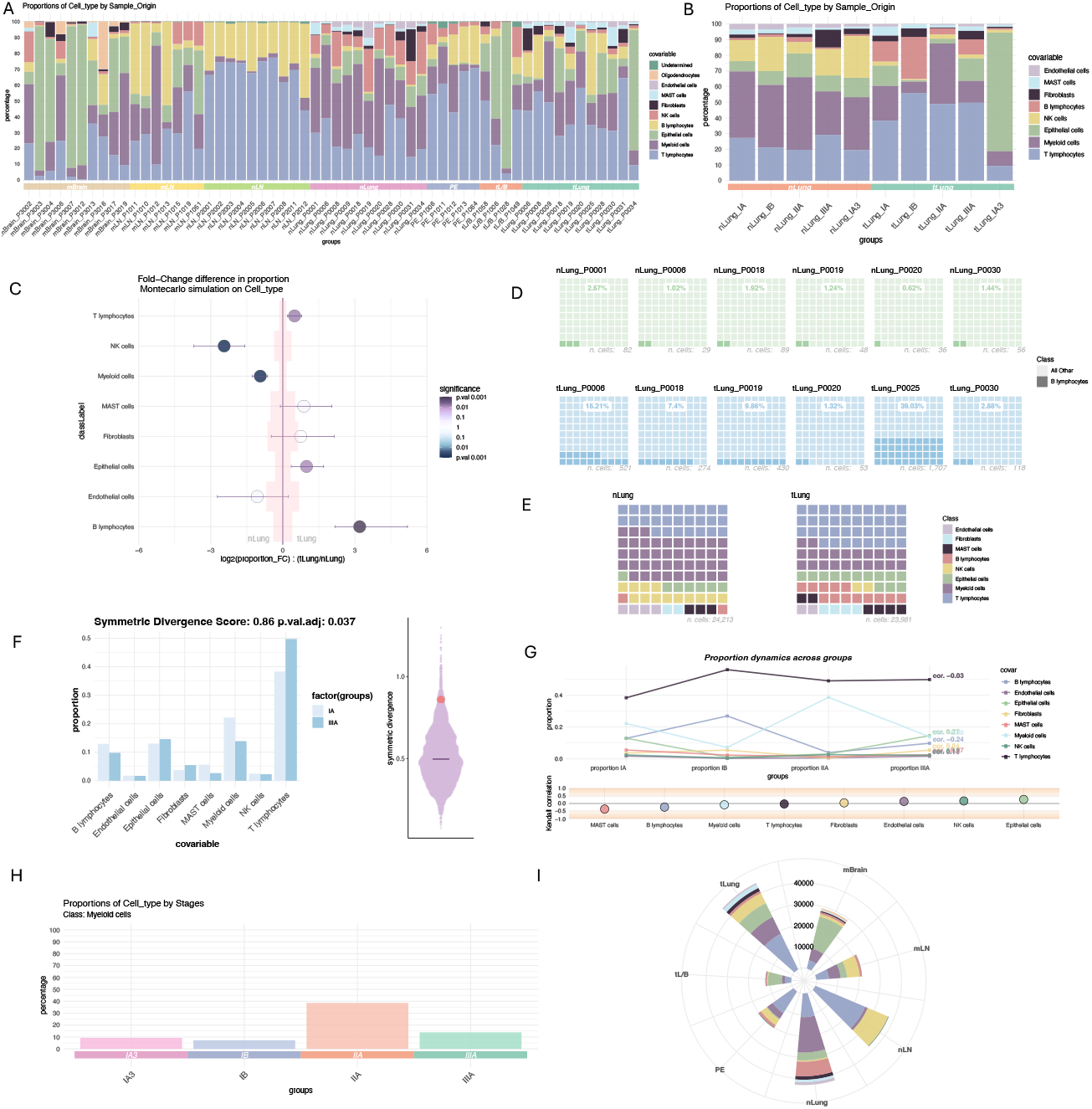
Diagram of plots and tests available in LotOfCells. A. Barplots of proportions of cell types for all individual samples from different tissues, the tissue class is depicted in the colored boxes on the x-axis. B. Barplots showing the cell type composition of normal and tumor lung samples, each bar corresponds to a cancer stage. Samples from the same stage-condition are grouped together in the same bar. C Montecarlo test for the difference in cell type population abundances between tumor and normal lung samples in stage IA. B lymphocyte population is significantly larger in tumor. D Waffle plots showing B lymphocytes only for all independent patients. The number of total B lymphocytes is depicted in grey. Each square=1%. E Waffle plot showing IA stage for tumor and normal lung. All samples are pulled together by condition. F Symmetric divergence score test between stages IA and IIIA from lung tumor. On the right a scatter violin plot showing the distribution of generated scores vs the observed value. G Cell type proportion changes across cancer stages in lung tumor. Values of the Kendall Tau correlation coefficient per cell type are shown. H Barplot of Myeloid cells proportions across cancer stages in lung tumor. I Polar plot showing the raw number of cells per tissue.

Here we show the capabilities of data visualization of our package, and more important, the agreement and significance assessment of the difference in proportions of relevant cell type populations that vary across cancer stages and conditions.

### 2.1 Cell heterogeneity in tumor and metastatic LUAD

We started by visualizing and comparing the major cell types populations from normal lung tissue (nLung) with the tumor Lung tissue (tLung, early stages). Patient heterogeneity was readily visible (Fig2B) both in normal and tumor lung samples. Differences in cell proportions found significant changes in both T and B lymphocyte (Fig2A), which were enriched in tumor lung (tLung), B lymphocytes in particular were extremely enriched (T lymphocytes: abundance FC: 0.67, p.value < 0.001; B lymphocytes: abundance FC: 2.99, p.value < 0.001), and a large decrease on the number of natural killer cells (NK abundance FC:-2.83, p.value < 0.001) and moderate decrease in myeloid cells (abundance FC: -0.99, p.value < 0.001) in tumor lung cancer (tLung) when contrasting the cell proportions to the normal lung samples (nLung). These results, in agreement with the original study [20], show the activation of the adaptive immune responses with T and B lymphocyte supporting major roles in the tumor micro environment. Additionally. we observed a small moderate increase in epithelial cells in the tumor lung tissue (abundance FC: 0.90, p.value < 0.001) (Fig2A). Additionally, to measure the heterogeneity of the normal lung samples (Fig2A), we used the symmetric divergence score from LotOfCells. First, we split the normal lung samples into two random groups, then we compute the score between the groups. As expected for normal tissue samples, we did not found a significant divergence score (p.value=0.303)(Fig2E), meaning that the visible heterogeneity per sample in normal tissues is not sufficient to incur in significant cell type composition differences.

### 2.2 Metastatic lymph node association with myeloid infiltration and progression

Visualization and comparison of cell subtypes in normal and metastatic lymph node tissues found a large heterogeneity in metastatic lymph node samples (mLN), and a much lower variability in normal lymph node (nLN) (Fig2C). When testing for cell abundance differences, we found a large number of significant cell sub-types more enriched in metastatic lymph node samples, including a large of immune cell types (Fig2D).

**Fig. 2:**
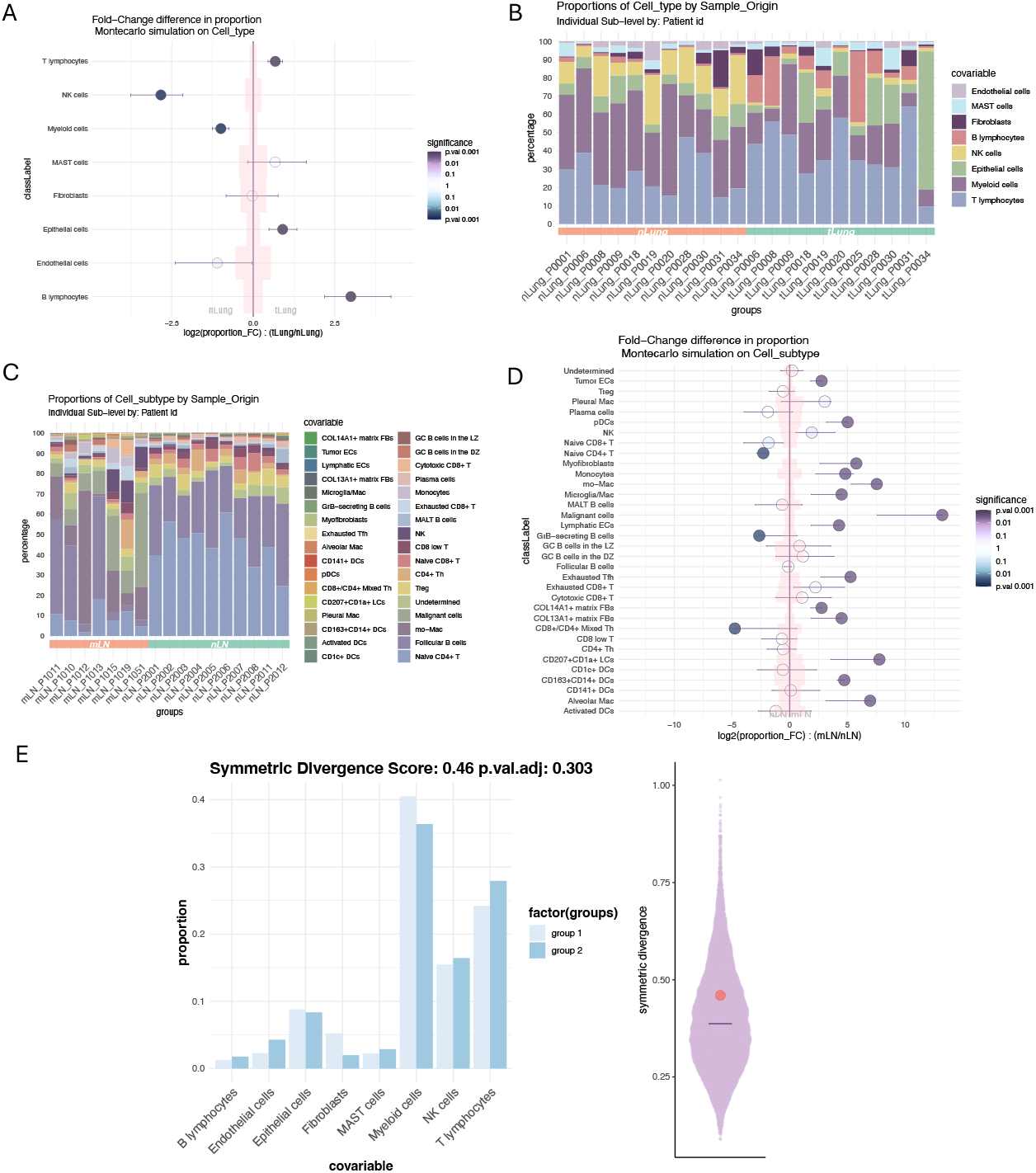
A. Differences in proportion on major cell types between normal lung (nLung) and tumor lung (tLung). B. Barplot showing major cell types per patient on normal and tumor lung. C. Barplot of cell subtypes across normal and metastatic lymph node samples (nLN, mLN). D Differences in proportions of cell subtypes across normal and metastatic lymph node samples (nLN, mLN). E. Symmetric divergence score between two random groups of normal lung tissue. The divergence is not significant, meaning that nor major differences in cell type abundances exists between normal tissue samples as expected.

In agreement with the original study [20], metastatic lymph nodes (mLN) had a significant larger number of myeloid cells in contrast with normal lymph nodes (nLN), indicating an association of myeloid infiltration with metastasis. Myofibroblasts were largely enriched in metastatic lymph node samples (abundance FC:5.76, p.value < 0.005), which supports a tumor progression role, specifically for fibroblastic reticular cells, which have been reported to be immunologically specialized myofibroblasts [20, 21].

### 2.3 Myofibroblasts and tumor progression

To analyse and visualise the distinct populations of fibroblasts, we selected this specific cell type from the metadata in to a subset. COL13A1+ and COL14A1+ matrix fibroblasts, pericytes, and Myofibroblasts showed a high degree of variation across tissues and stages (Fig.2A,B). To test whether any cell type proportion changes significantly across different lung cancer stages (nLung:normal lung, tLung: tumor Lung and tL/B: tumor Lung advance stage), we used LotOfCells to compute the Kendall correlation coefficient (see Materials and Methods). Results showed (Fig.2C) that myofibroblasts (Kendall Tau:0.97, p.value < 0.001) and COL14A1+ matrix fibroblasts (kendall Tau: -0.98, p.value < 0.001) were the only significant cell types to vary in proportions across stages, in accordance with the original study [20], supporting the shift of fibroblasts towards promoting tissue remodeling and angiogenesis in LUAD, specifically Myofibroblasts, which are know to promote tissue remodeling, angiogenesis, and tumor progression [22–24].

## 3 Discussion

Here we show how to statistically assess whether the proportion of cell numbers vary significantly between conditions, or even across a series of stages. We believe that this tool will help to clarify and easily test quantitative differences that usually are claimed on single cell studies albeit commonly not statistically tested.

**Fig. 3:**
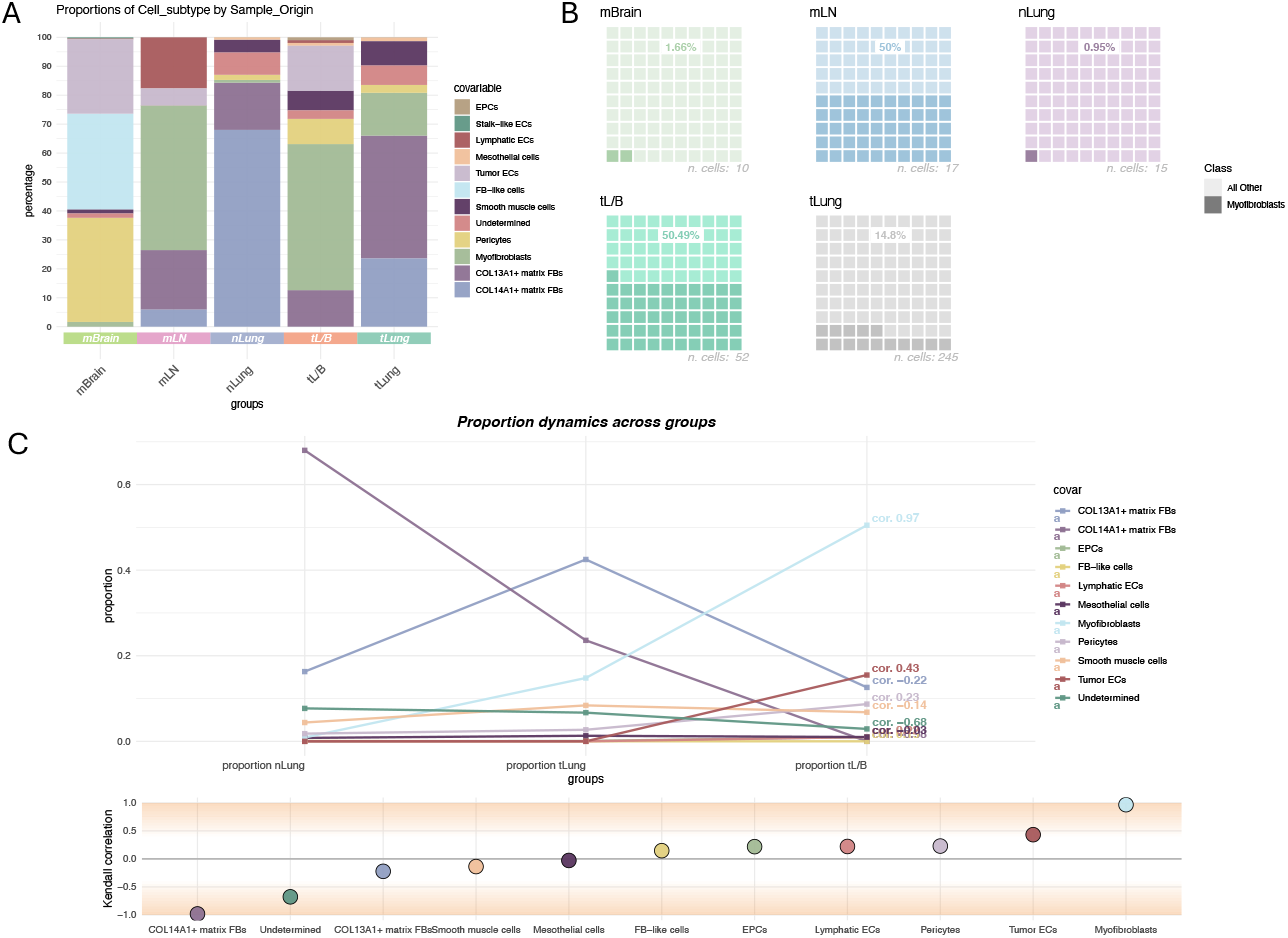
A. Barplot showing major cell types in fibroblast subset across tissues. B. Waffle plot showing myofibroblasts distributions across tissues. C. Kendall correlation test measuring differences in cell type abundances across normal, tumor and advance lung tissue (nLung, tLung, tL/B)

## 4 Materials and Methods

### 4.1 Montecarlo simulation for population frequency differential analysis

#### 4.1.1 Two-condition comparison

In order to compute a null distribution in each iteration, we randomly sample *sqrt* (*n*_*ik*_) cells from each *class*_*i*_ - *condition*_*k*_, then we compute the frequencies that each *class*_*i*_ represents over the randomly sampled population *sqrt* (*n*_*ik*_) on *condition*_*k*_. If a sample id (or additional sub-level) is defined, then sub-sampling *sqrt* (*n*_*ijk*_) is perform individually for each *class*_*i*_ *sample*_*j*_ *condition*_*k*_ to account for sample cell number variability, then sampled cells are pulled together for each *class*_*i*_ and frequencies are computed.

Before calculating the frequencies and to deal with zero cell cases, we add a *pseudo-count* based on the hyperbolic arcsine (asinh) function with no log transformation:

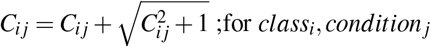

Finally, we compute the frequency fold change (FC) for each class *i*:

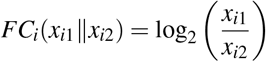

#### 4.1.2 Comparison of more than two ordered-conditions

If more than two conditions are defined, a correlation test using Kendall’s Tau-a will be perform to test whether the increase or decrease in proportions per class correlates with the order of the conditions given. The random sub-sampling is perform as described in the previous section.

The Kendall Tau correlation coefficient is a measure of correlation between two ranked variables. It is defined as:

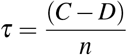

where:

- *C* is the number of concordant pairs.
- *D* is the number of discordant pairs.
- *n* is the total number of pairs, given by 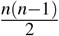.

### 4.2 Symmetric Divergence Score

Our proposed method introduces a symmetric score, based on the Kullback-Leibler (KL) divergence as a measure of global dissimilarity in class distributions between two samples (e.g.: global cell-type distribution shifts in scRNA-seq data due to a covariate). The symmetric score is based on the Kullback-Leibler (KL) divergence, a measure of relative entropy between probability distributions.

Given two samples with proportions *x*_*i*1_ and *x*_*i*2_ for each class *i* (e.g., cell types), the KL divergence 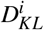 for each class is calculated as:

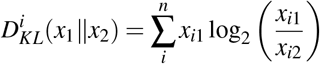

The symmetric score is computed by summing the KL scores from both scenarios:

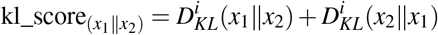

For the significance test we again, as described in previous sections, do a permutation analysis in which we randomly pull cells to construct a null distribution of symmetric divergence scores, then we compare our observer value to the distribution of generated scores. Beeswarm violin plot of the null distribution values is build using the ggbeeswarm R package [25].

### 4.3 Significance testing

For significance testing we recommend and *α* value of 0.001 for all the test described and included on the package LotOfCells.

## 5 Code availability

The r code package, as well as examples and the manual, can be accessed on the GitHub repository: https://github.com/OscarGVelasco/LotOfCells

## Acknowledgments

This publication was supported through state funds approved by the State Parliament of Baden-Württemberg for the Innovation Campus Health + Life Science Alliance Heidelberg Mannheim.

